# nearBynding: A flexible pipeline characterizing protein binding to local RNA structure

**DOI:** 10.1101/2020.10.24.352591

**Authors:** Veronica F. Busa, Alexander V. Favorov, Elana J. Fertig, Anthony K. L. Leung

## Abstract

The etiology of diseases driven by dysregulated mRNA metabolism can be elucidated by characterizing the responsible RNA-binding proteins (RBPs). Although characterizations of RBPs have been mainly focused on their binding sequences, not much has been investigated about their preferences for RNA structures. We present nearBynding, an R/Bioconductor pipeline that incorporates RBP binding sites and RNA structure information to discern structural binding preferences for an RBP. nearBynding visualizes RNA structure at and proximal to sites of RBP binding transcriptome-wide, analyzes CLIP-seq data without peak-calling, and provides a flexible scaffold to study RBP binding preferences relative to diverse RNA structure data types.

## Background

RNA-binding proteins (RBP) play diverse, important roles from RNA biogenesis to degradation. Over 1,500 human genes code for RBPs, making them among the largest families of the human proteome [1]. Dysregulation of RBP binding can drive diseases such as cancer, muscular dystrophy, neurodegeneration, and developmental abnormalities [2–4]. Parsing RBP binding specificity and function is therefore central in understanding human biology and diseases. Most RBPs demonstrate at least some RNA sequence, structure, or modification preferences in their binding sites [5–7]. Some RBPs recognize RNA structure more than sequence [8,9], but binding preferences to structured RNA have thoroughly been described for only a few proteins, and RNA structure surrounding protein binding events is rarely characterized.

One of the most precise methods for determining transcriptome-wide RBP binding is UV cross-linking immunoprecipitation with deep sequencing (CLIP-seq). Current practice when analyzing CLIP data to identify RBP binding sites is to call peaks, which binarizes data to intervals of bound or unbound RNA. Although peak-calling simplifies many downstream analyses, this binarization sacrifices information that could be gleaned from relative peak intensity and amplifies the error at miscalled positions. Analyzing CLIP data in a non-binarized format potentially enables more nuanced motif-finding by preserving levels of binding preferences in the input data. Motifs ascribed to RBPs are often insufficient for explaining a large proportion of binding occurrences [10–15]. Describing the unexplained binding of RBPs—especially for RBPs that bind structured RNA—will increase our potential to elucidate the etiology of diseases driven by dysregulated mRNA metabolism.

Besides sequence, RBP binding can also be impacted by RNA structures. Therefore, integration of nucleotide-resolution RNA folding information with CLIP-seq is critical to discern transcriptome-wide RBP binding. Unfortunately, RNA structures are diverse and much more difficult to discern than RNA sequence. Currently, high-throughput RNA structure is obtained using *in silico* prediction algorithms such as Sfold, RNAfold, or RNAshapes [16–18]. In their simplest form, RNA structure prediction algorithms input the sequence of the RNA and discern probable RNA structure using pre-coded, experimentally-defined biophysical metrics. Sequence-based RNA structure predictions are, however, unable to account for how RNA modifications or RBPs may impact RNA structure *in vivo* because they calculate folding of a naked RNA strand. Despite these assumptions, predicted structure information provides a valuable complement to CLIP-seq data as it provides an approximation of the geometry of the RNA landscape where the RBP binds.

Several machine learning algorithms have been developed to resolve structure-based RBP motifs using CLIP data and RNA structure prediction [19–21]. Binding occurrences of some RBPs are almost accurately predicted by some of these machine learning algorithms such as RPI-Net. For example, selected datasets of TAF15, ELAVL1, and HNRNPC have area under the receiver operating characteristic [AUROC] >0.98 (where 1 is perfect classification and 0.5 is random chance), but the binding preferences of others remain difficult to predict with these approaches (e.g. AUROC = 0.724 for ALKBH5 and <0.85 for ZC3H7B, C22ORF28, and C17ORF85 datasets), and many datasets for RBPs have not been explored [21]. The dampened prediction accuracy of these RBPs is perhaps due to biological confounders, such as subcellular localization or cofactors that affect binding preference, which cannot be discerned from CLIP data alone. The predictive power of these state-of-the-art algorithms may be limited by their reliance exclusively on sequence-based RNA structure prediction and their lack of accommodation for experimentally-derived RNA structure information (reviewed [22,23]). A further limitation of these algorithms is that they predict RNA structures only from short intervals [22]. Although more computationally efficient, this approach sacrifices the possibility of long-range contacts that affect secondary structure or of tertiary conformations, which may shield or expose binding sites.

Visualization techniques serve as another critical complement to interpreting binding predictions. Algorithms such as GraphProt and iDeepS incorporate a post-processing step to easily visualize the sequence and structure preferences [19,20], but these algorithms only provide visualization of structure information for a short binding motif (7-12 nucleotides). They also do not offer insight about the RNA structure surrounding those motifs, despite evidence that such local context can be important in RBP binding [24,25]. On the other hand, current methods interrogating RNA structure binding contexts beyond the binding site can only be performed in a low-throughput manner [24,25]; therefore, RBPs known to have preferred proximal secondary structures are sparse.

We develop a new algorithm nearBynding to model RBP binding through the integration of CLIP-seq and RNA structure data, which can be derived from *in silico* predictions or experiments. The nearBynding pipeline is unique in three ways: first, it visualizes RNA structure at and also proximal to RBP binding sites in a transcriptome-wide manner; second, it is a flexible scaffold to study RBP binding preferences relative to diverse RNA structure data types; and third, it can analyze RBP binding from CLIP-seq data without peak-calling. We perform this integration by extending the kernel correlation algorithm StereoGene [26] to assess correlation between continuous or interval features from RNA structure and RBP binding along the transcriptome. Directly estimating correlation among pairs of continuous or interval features allows users to analyze track features associated with binding and RNA structure without peak calling and associated data loss. Notably, the cross-correlation model in StereoGene can also represent trends between proximal coordinates of RNA structure and CLIP-seq to allow for identification of extended binding structure. We benchmark nearBynding using simulated data and replicate CLIP experiments. We demonstrate its utility by comparing our results to known RBP binding preferences, employing diverse data types to predict RBP binding preferences that are unusable by currently-available RBP motif-finding software, and using these discovered RBP binding preferences to hypothesize RBP characteristics that may predispose binding preferences for or against specific RNA structures.

The software for nearBynding is developed in an R/Bioconductor package by the same name to enable broad usage. In its simplest form, the nearBynding software estimates RNA binding preference from a BAM file of aligned CLIP-seq reads and a list of transcripts that correspond to an annotated genome in GTF format. The software also allows extensions for additional inputs, such as experimental RNA annotations including nucleotide modification data to complement RBP binding. This additional integrative analysis leverages both data sources to offer insights into RNA structure that nucleotide sequence alone cannot provide and makes this pipeline a diverse, flexible tool to study the context of RBP binding.

## Results

### Overview of the nearBynding pipeline for transcriptome-wide RBP binding prediction

We present nearBynding—a pipeline that incorporates RBP binding sites and RNA structure information to discern local RNA structure for regions bound by an RBP (Fig. 1A). To visualize the local binding context of an RBP, nearBynding uses a list of transcripts, an annotated genome, and aligned CLIP-seq data as inputs. RNA structure and RBP binding data is mapped to a concatenated transcriptome produced from only the transcript intervals of interest, and StereoGene [26] is called to calculate the cross-correlation of RBP binding from CLIP-seq data and RNA structure.

**Figure 1.**
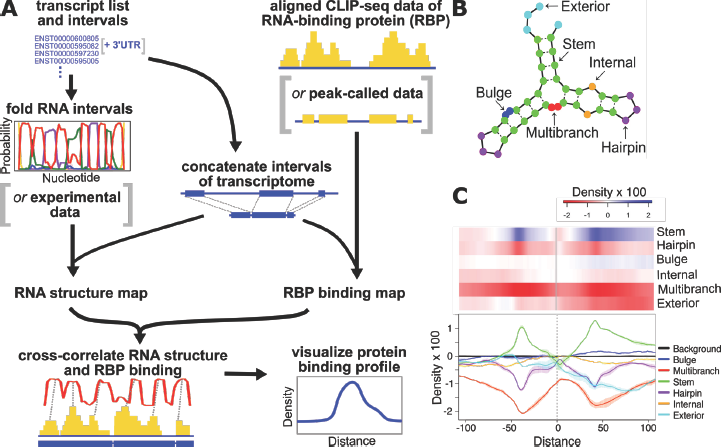
Overview of the pipeline. **A.** The user inputs CLIP-seq data (aligned reads or called peaks), a list of transcripts, and an annotated genome. Optionally, *in silico* folding of RNA intervals can be replaced by an experimentally-derived RNA structure track. RBP binding and RNA structure data are mapped to the concatenated transcriptome and cross-correlated. The nearBynding pipeline outputs cross-correlation densities and their distributions to estimate RNA binding. **B.** Examples of six RNA structure contexts predicted by CapR [6] for which the nearBynding pipeline can be applied. **C.** Example heatmap and line plot visualizations of PUM2 binding from eCLIP data in two replicates of K562 cells estimated from the cross-correlation densities and visualized as part of the nearBynding software. The line plot shows the average signal as a dark line and error bars as a lighter-colored shading. The heat map only shows the average value at every position if multiple samples are used to calculate density values to model binding.

RNA structure contexts can be categorized into one double-stranded context (stem) and five single-stranded contexts: hairpin, multibranch, internal, exterior, and bulge (Fig. 1B). By default, the nearBynding pipeline uses RNA structure probabilities predicted from sequence by CapR [6] for the selected transcriptomic intervals (see Methods). While CapR provides the default structural data input, the computational methods in the pipeline can be adapted for alternative inputs of custom RNA structure tracks or intervals, such as RNA modifications that affect RNA structure (e.g. N^6^-methyladenosine [m6A]). In all cases, StereoGene generates cross-correlation densities for all RNA folding contexts relative to RBP binding. Since cross-correlation shows the relative position of one track (e.g., RBP binding) to another track (e.g., RNA structure), we can use it as a tool to visually represent RNA structure context upstream, at, and downstream of RBP binding.

Binding profiles illustrating RNA structures at and proximal to RBP binding can be visualized either as line plots with standard errors for cases with multiple replicates or as heatmaps (Fig. 1C). These simple visuals provide a holistic RBP binding profile that can be used to predict RBP binding transcriptome-wide, especially in cases where mutations may modify RNA structure. In addition to allowing visual assessment, the nearBynding pipeline includes functions to quantitatively compare RBP binding motif cross-correlation distributions between two different RBPs. Specifically, users are able to pairwise compare the Wasserstein, or earth-mover, distance [27] between RBP binding profiles. For example, a short Wasserstein distance suggests similarity between two RBP profiles, which may imply binding competition or cooperation between RBPs.

In the default analysis pipeline implemented in the nearBynding software, CLIP-seq samples can be input as a BAM file containing aligned reads or as a BED file containing peak intervals or protein-RNA cross-linked sites. Some input data, such as binding peaks or cross-linking sites identified by other programs, have already corrected the data for background signals. However, in cases where the input is raw aligned CLIP data, it is prudent to provide a background input to the nearBynding pipeline to ensure the observed signal is from the RBP of interest rather than from experimental artifacts such as cell-specific transcript levels or size-matched input noise. For example, the binding profiles for HNRNPK in HepG2 and K562 cells were much more similar after background signal was removed (Wasserstein distance of 1.80 between non-corrected profiles versus 0.23 for background-corrected profiles) (Fig. S1).

### Benchmarking on simulated data

The nearBynding pipeline is unique because it can visualize RNA structure at and proximal to RBP binding, incorporate diverse RNA structure data types, and analyze minimally processed CLIP-seq data. Because of the uniqueness of this pipeline, we were unable to directly compare it to other algorithms available, so we designed simulated data studies to benchmark a full range of biological variables that may impact performance (Fig. 2). Briefly, we tested three factors that may impact RBP binding context signal strength: peak concordance between tracks, foreground to background ratio of the RBP binding track, and peak width range of the RBP binding track. Each simulation contained a pairwise analysis of the cross-correlation between an RNA structure track [RNA] and a CLIP-seq track [CLIP], where a greater amplitude for cross-correlation density reflects better co-occurrence of the two tracks. The RNA structure track peak distances and heights were varied to simulate the range of predicted RNA structure probabilities and random distribution of these structures across the transcriptome. The RNA structure track consisted of 10,000 peaks 31 to 500 units apart (unless otherwise stated), 5 units in width, and 0.02 to 1 units in height. The CLIP-seq track simulated signal from aligned CLIP-seq data and contained a mixture of both background and foreground signal. Since the width of RBP binding peaks cannot be narrower than the length of CLIP-seq sequencing reads, the CLIP-seq track contained 30-unit-wide peaks (unless otherwise stated) to simulate the 30-nt reads of CLIP-seq data deposited in the ENCODE portal [28]. The CLIP-seq track was also shifted 12 units to the left of and equal in height to the RNA structure track peaks.

**Figure 2.**
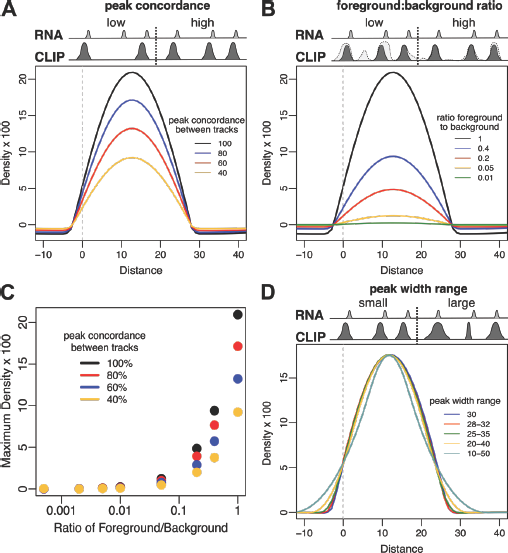
Cross-correlation distribution tracks of simulated RNA binding data to benchmark the performance of the nearBynding pipeline. In the simulations, the RNA structure track [RNA] is shifted twelve units to the right relative to the CLIP track [CLIP] to model proximal RNA structure, as reflected by the maximum cross-correlation density at distance=12. For A, B, and D, the middle grey peaks represent RNA structure data and the dark grey peaks represent CLIP simulation data. **A**. Cross-correlation distribution tracks with differing peak concordance. **B.** Cross-correlation distribution tracks with differing foreground to background signal ratios. The lightest grey regions of peaks represent background. **C.** Maximum cross-correlation density values for pairs of tracks with varying peak concordance and foreground to background signal ratios. **D.** Cross-correlation distribution tracks with differing peak width range.

First, we tested the impact of the frequency of an RBP binding to its target RNA structure across the transcriptome, which may be affected by the accessibility of the RNA structure and the binding strength of the protein. To simulate this effect, we varied the frequency of the foreground signal peak concordance of the simulated CLIP-seq track relative to the simulated RNA structure track (Fig. 2A). We hypothesized that tracks with a higher frequency of RBP binding to target RNA structure would provide stronger binding signals than RBPs with sparser target binding. Supporting this hypothesis, the result of the nearBynding pipeline for the simulated data showed that cross-correlation signal strength correlates positively with peak concordance.

Next, we simulated artifacts associated with collecting CLIP-seq reads, such as background signal from input by varying the height of the background signal of the simulated CLIP-seq track relative to the foreground signal (Fig. 2B). We hypothesized that simulations with a greater foreground (dark grey) to background signal (light grey) would have stronger RBP binding signals. As expected, cross-correlation signal strength correlated positively with the ratio of foreground to background. Both peak concordance and foreground to background ratio greatly affected signal strength, with the nearBynding pipeline requiring a foreground to background signal greater than 0.05 to detect the binding signal (Fig. 2C). Therefore, our pipeline performance may be optimal when applied to data collected by protocols that minimize noise (e.g., via additional washing steps) rather than protocols that document all binding events at the expense of greater noise.

We further employed our simulated data to test the sensitivity of nearBynding to the uniformity of binding peak width. Specifically, we increased the range of the simulated CLIP-seq peak widths to accommodate the possibility that RBPs may have variable binding footprints (Fig. 2D). Though the shape of the cross-correlation density track changed to reflect greater variation in peak widths, the amplitude and position of the signal maximum did not. Therefore, we conclude that differences in peak width have no effect on signal amplitude.

Our pipeline relies on a concatenated transcriptome as an input for StereoGene. However, RBP-bound transcripts may not be evenly distributed across the concatenated transcriptome. Therefore, we tested the dependence of nearBynding on the distribution of peaks along a concatenated transcriptome by shifting the locations of the simulated peaks such that they were all uniformly distributed or clustered near either end of the CLIP-seq track (Fig. S2). Compared to peak concordance and foreground to background ratio, only a negligible loss in signal amplitude was observed for the most extremely skewed data (Fig. S2). Overall, our results demonstrate that the order in which transcripts are concatenated, which could possibly affect the distribution of peaks, has negligible effect on binding signal relative to other variables tested.

### Cross-correlation tracks reproducibly cluster RBP data across biological replicates

The context-dependence of RNA binding can be expected to lead to variable signal concordance in binding predictions from CLIP-seq data with the same RBP. Replicates from the same cell type would likely manifest technical differences whereas analyses of the same RBP across different cell types may depict biological differences in RBP binding. Analyses of the same RBP between replicates within the same cell type can be expected to have greater concordance than analyses from different cell types. We sought to make qualitative assessments about the fidelity of the nearBynding pipeline’s ability to reproducibly identify such RBP binding context by clustering RBP binding profiles.

The ENCODE portal [28] has enhanced CLIP (eCLIP) datasets for 103 RBPs in HepG2 and 120 RBPs in K562, with each dataset containing two replicates and an input control. Of these, 73 RBPs are common across both cell lines. Genome-wide RNA structure profiling showed that 3’ untranslated regions (UTR), which are targets for many RBPs, are generally highly structured in cells [29,30]. Therefore, in order to test our pipeline on a robust dataset, we restricted our analysis of RBP binding to 3’UTR regions. We collected isoform information of all 3’UTRs expressed in HepG2 and K562 using RNA-seq data from ENCODE [28]. We generated cell type-specific binding profiles by selecting eCLIP reads that aligned to isoforms expressed in the corresponding eCLIP cell type. The 3’UTRs for the expressed transcripts were then submitted to the nearBynding pipeline to determine RBP binding preferences for these regions.

First, we wanted to test how well biological replicates of the same RBP in the same cell type clustered. We used the Wasserstein distance [27] to determine the amplitude and distance required to transform one RBP binding profile into another. We calculated the sum of Wasserstein distances between the cross-correlation density tracks of all RNA structure contexts for every sample within each cell type. 71 of 206 replicates in HepG2 (34%) and 115 of 240 replicates in K562 (48%) most closely clustered in pairs with their corresponding biological replicate, and 50% of HepG2 and 58% of K562 replicates were within the top three closest distances of their corresponding biological replicate (Fig. 3A and S3A).

**Figure 3.**
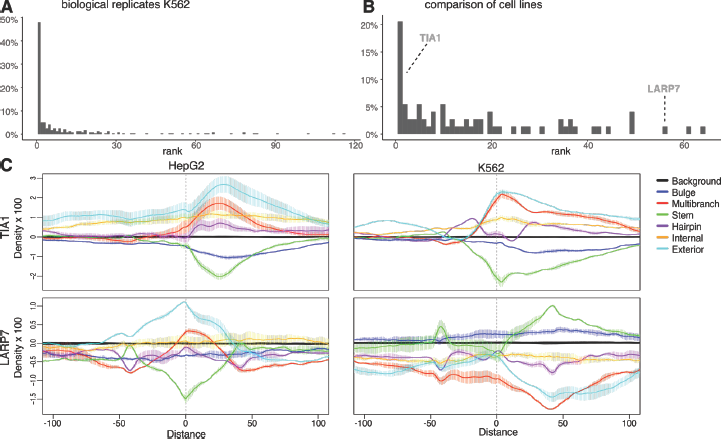
Binding profiles of all RBPs with eCLIP data in ENCODE clustered using Wasserstein distance. **A.** Histogram of ranks for Wasserstein distances of paired biological replicates in K562 cells. **B.** Histogram of Wasserstein distance ranks for the same RBP across HepG2 and K562 cell lines. The ranks of TIA1 and LARP7 across cell lines are indicated. **C.** Example binding profiles for TIA1 (top), an RBP that is similar across cell types, and LARP7 (bottom), an RBP that is dissimilar across cell types, in HepG2 (left) and K562 (right) cells.

Next, we interrogated the reproducibility of RBP binding profiles across cell lines. The cross-correlation densities of biological replicates for each RBP were averaged, and these averaged values were used to calculate the Wasserstein distances for all RNA structural contexts. For every RBP in K562 cells, we ranked how similar its binding profile was to RBPs in HepG2 cells. 15 of 73 RBPs (21%) clustered closest with their counterparts in the other cell line, and 21 of 73 RBP counterparts (29%) were within the top three closest distances in the other cell line (Fig. 3B). The inverse comparison—the distance of HepG2 RBPs against all K562 RBPs—also had 29% of RBPs cluster within the top three distances of their counterparts (Fig. S3B). The concordance of RBP binding profiles across cell lines were lower than biological replicates of the same cell lines. This difference may be either due to the distance algorithm disproportionately weighing noisy signals and placing dissimilar binding profiles close and similar profiles distant from one another or an RBP having dissimilar binding profiles between cell types.

To distinguish between these possibilities for why not all RBPs cluster closely across cell lines, we visually inspected binding profiles to test whether qualitative assessments align with the Wasserstein distance algorithm. We randomly selected 25 binding profiles that clustering labelled as either similar (rank 3 or less; e.g., TIA1) or dissimilar (rank 30 or greater; e.g., LARP7) across cell types (Fig. 3C and Table S1). We then performed crowd-sourced validation, where ten individuals were asked to visually bin these same pairs as either similar or dissimilar based on a 200-nucleotide binding profile window. All 25 pairs tested were binned accurately by the majority of validations (Fig. S3C), demonstrating that Wasserstein distance is a reliable means of identifying similar samples. We concluded that RBPs with binding profiles across cell types that cluster tightly have similar binding profile shapes, whereas RBPs that do not cluster across cell types have dissimilar binding profile shapes.

### RBP binding profiles recapitulate known structural preferences

Next, we tested whether the binding profiles generated by nearBynding reflect known RBP structural binding preferences. Although PUM2 preferentially binds 3’UTRs in a sequence-specific manner, there is evidence that PUM2 also has a structural component to its binding preferences: *in vitro* analysis shows that PUM2 dissociates from double-stranded regions faster than single-stranded regions and that it stably binds regions flanked by stem structures [25]. The PUM2 binding profile (Fig. 1C) showed that PUM2 has minimal structure preference at the point of binding (distance = 0), but it does prefer stem context upstream and downstream of its point of contact.

PUS1 has a weak trinucleotide binding sequence motif and modifies nucleotides at the 5’ end of stem loops flanked by single-stranded runs for the vast majority of its high-confidence targets [24]. Consistent with PUS1 binding and modifying the 5’ base of stems, its binding profile showed a preference for single-stranded regions at the end of the transcript (exterior context) upstream and double-stranded (stem) context downstream of PUS1 binding (Fig. 4A).

**Figure 4.**
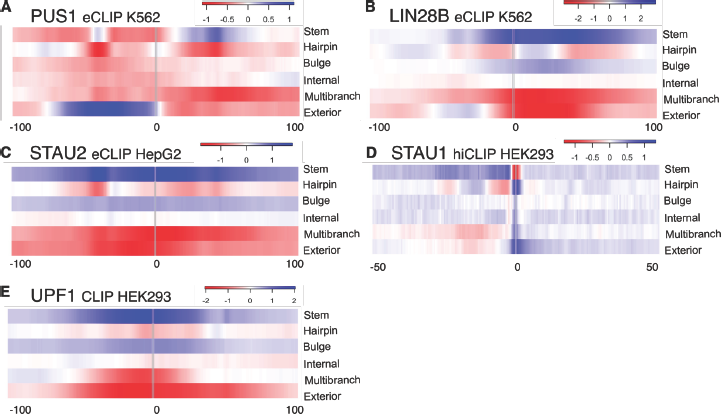
Application of the nearBynding pipeline to analyze RBP binding profiles for proteins with known structure preference. Binding profiles for **(A)** PUS1, **(B)** LIN28B, and **(C)** STAU2 from eCLIP data. **D.** Binding profile for STAU1 from hiCLIP cross-link site data. **E.** Binding profile of helicase-dependent UPF1 binding based on subtraction of DEAA UPF1 signal from WT.

LIN28B has two RNA-binding domains: a cold shock domain (CSD) and tandem zinc-binding motifs (ZFs). Although LIN28 has a preference for binding GGAGA motifs, target motifs are generally single-stranded [13]. NMR spectroscopy suggests that although LIN28 binds stem-rich regions, the CSD binds hairpins and the ZFs bind bulges containing the sequence motif associated with the stem [31]. These same results were apparent in the binding profile, which showed enrichment for stem, bulge, and hairpin contexts at or proximal to the LIN28 binding site (Fig. 4B).

STAU2 contains three double-stranded RNA-binding domains (dsRBDs) and binds stretches of base-paired sequences of variable lengths [32]. Although the dsRBDs bind tightest to perfectly complementary stem structures, they are able to bind stems that contain bulges [32]. Consistent with expectations, the binding profile of STAU2 was strongly enriched for stem context, had slight enrichment for bulge context, and was generally depleted for single-stranded contexts such as hairpin, multibranch, and exterior (Fig. 4C).

There is a range of resolutions among CLIP techniques. For example, eCLIP provides 30-nucleotide reads surrounding the protein–RNA cross-linking site, whereas better resolution can be achieved with techniques such as individual-nucleotide resolution CLIP (iCLIP) and RNA hybrid iCLIP (hiCLIP) that are able to identify the RNA–protein cross-link site with single-nucleotide resolution. The resolution of nearBynding’s profiles reflects the resolution of the input data. For example, by using hiCLIP cross-link sites of STAU1 [33], which binds dsRNA similar to STAU2, nearBynding was able to demonstrate that STAU1 contacts ssRNA—preferably hairpin context and possibly other ssRNA contexts at the binding point—but that this ssRNA was directly 3’ of the double-stranded stem context (Fig. 4D). The authors of the hiCLIP data hypothesized that cross-linking sites were enriched at ssRNA because bases within the duplexes are inaccessible for protein–RNA cross-linking [33]. Further, the cross-linking site was often 3’ of a stem–hairpin–stem structure. Although there are only a few experimentally-confirmed RNA structure binding preferences for us to use as true positives, our binding profiles effectively recapitulate documented RBP preferences.

Besides investigating wild-type (WT) protein binding relative to background signal, nearBynding can be applied to researching mutant RBPs by comparing WT and mutant protein binding. Whereas a comparison of WT versus input control depicts the full complement of RBP binding across the transcriptome, a comparison of WT versus a mutant allows visualization of the function-dependent binding of an RBP. For example, binding data is available for WT UPF1 as well as two helicase-dead mutants, K498A and DEAA, which are deficient in ATP binding and hydrolysis, respectively [34]. Both helicase-dead UPF1 mutants retain the ability to bind RNA, but they exhibit a complete loss in target discrimination [34]. The binding profiles of WT UPF1 minus helicase-dead signals suggested that UPF1 requires helicase activity to occupy stem contexts and select against the unstructured multibranch and exterior contexts (Fig. 4E and S4).

### Called peaks or aligned tracks for RBP binding produce similar binding profiles

Current practice for analyzing CLIP data is to call RBP-bound peaks using algorithms such as CLIPper [35] and Piranha [36]. We selected 29 different RBPs in HepG2 and K562 cells that demonstrate strong, reproducible binding signals at 3’UTRs based on analysis from ref. [30] (Fig. S5A), which came to 40 unique cell type–RBP combinations. We used these high-confidence datasets to test whether nearBynding can produce comparable peak binding profiles from peak-callers and aligned reads. We collected eCLIP aligned reads for these RBPs from the ENCODE portal [28] and ran Piranha on all replicates with parameters as described in the original paper [36]. We also downloaded CLIPper-derived peaks of the eCLIP data from the ENCODE data portal [35]. These three different inputs—aligned eCLIP tracks, Piranha peaks, and CLIPper peaks—were run through the nearBynding pipeline, and binding was assessed for 3’UTR-annotated regions of the transcriptome. We calculated Wasserstein distances between all 40 unique cell type–RBP combinations with 3 different input types each. Visualizing their distances in 2D on a multidimensional scaling plot showed only minor differences in the binding profile for RBPs based on the input source (Fig. S5B). Bootstrap analysis further indicated that the binding profiles for the three input sources of the same protein are closely clustered compared to randomly chosen binding profiles from other proteins in the same cell line (Fig. S5C). Therefore, the difference between profiles for different RBPs is far greater than the difference within the same experiment queried via different inputs.

### Inform RBP binding preferences using experimentally-derived RNA annotations

nearBynding is not restricted to *in silico* RNA structure prediction input, so we next interrogated RBP binding profiles with experimentally-derived RNA structure data. Guanine-rich RNA sequences can interact via Hoogsteen base-pairing and fold into non-canonical structural motifs called Gquadruplexes (G4s) [37]. Although many tools are available to predict putative G4s, they are prone to false-positives, since G4 folding is often dependent on the wider context of the RNA sequence and RBP regulation [29,38]. We therefore used rG4-seq data [39] to map G4s that form in cells. Although the rG4-seq data was collected from HEK293 cells and ENCODE provided RBP binding data from HepG2 and K562 cells, we reasoned that these cell lines would have enough G4s in common that we could observe general G4 binding trends. Indeed, we observed strong RBP binding at G4s for multiple published G4-binding proteins such as NONO, FUS, GRSF1, DROSHA, and DDX42 (Fig. 5A and S6A) [40–44]. Additionally, many of the RBPs that exhibited the strongest G4-binding signal—PRPF4, GTF2F1, FAM120A, CSTF2T, and DDX6—have recently been shown to bind at putative G4 sites in mRNA UTRs [45]. However, some published G4-binding proteins such as FMR1 did not exhibit a robust signal, perhaps due to cell-type-specific variations in binding (Fig. 5A and Table S2). Our analysis also identified RBPs such as YBX3, PRPF8, ZNF800, PPIG, and NOLC1 that are depleted for G4s at their binding sites in HepG2 and K562. However, these proteins have not previously been documented for their preference against G4 binding, which warrants future investigation.

**Figure 5.**
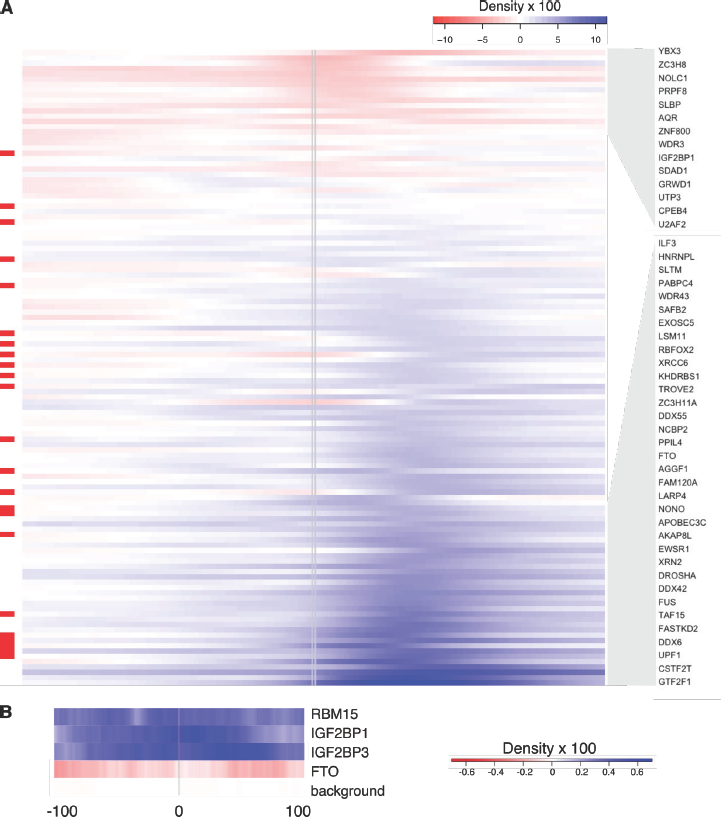
**A.** G4 binding profiles for all 120 proteins with eCLIP data in K562 cells. RBPs with molecular evidence of G4 binding in the literature are indicated in red on the left. RBPs with the most positive and most negative correlation signals are highlighted by the grey blocks and listed on the right. **B.** m6A binding profiles for RBM15, IGF2BP1, IGF2BP3, and FTO based on miCLIP-seq data.

We wondered whether our protein-level data can help identify domains that play a role in G4 binding preference. To increase the amount of data available, we pooled HepG2 and K562 data and took the maximum G4 density signal—positive or negative—from every RBP’s binding profile as a metric for G4 binding preference. We used the Pfam database [46] and protein sequence information to identify protein domains present within the RBPs. Across 13 common protein domains identified, most did not affect G4 binding (Table 1, Fig. S6B). RGG repeats are the most common motif in G4-binding RBPs (e.g., FUS) [47] and, based on our analysis, RBPs with RG-rich domains did demonstrate increased G4 binding. Proteins that contain SAP, dsRBD, and G-patch domains also had increased G4 binding, though there is no literature evidence of this preference. In contrast, RBPs that contain an armadillo domain had significantly decreased G4 binding, with 6 out of 8 armadillo-containing proteins demonstrating G4 depletion in their binding preference.

**Table 1.** Influence of RBP domains on G4 binding signal. Statistics from pooled HepG2 and K562 binding profiles. G4 signal difference is the difference in mean G4 binding signal between proteins with and without the indicated domain. Effect size is Cohen’s *d* and the p-value is of a Welch’s t-test comparing G4 signal of proteins with and without the indicated domain.

| Domain | G4 Signal Difference | Effect Size | p-value |
| --- | --- | --- | --- |
| armadillo | -2.41 | 0.913 | 0.00147 |
| RGG-repeat | 1.57 | 0.592 | 0.00612 |
| SAP | 0.836 | 0.312 | 0.0123 |
| dsRBD | 1.57 | 0.589 | 0.0219 |
| G-patch | 1.77 | 0.662 | 0.0385 |
| alpha-beta | 0.476 | 0.178 | 0.198 |
| winged-helix | 1.88 | 0.707 | 0.238 |
| RRM | 0.385 | 0.144 | 0.301 |
| K-homology | -0.384 | 0.143 | 0.402 |
| helicase | 0.428 | 0.16 | 0.436 |
| WD40 | -0.479 | 0.179 | 0.66 |
| P-loop | 0.0385 | 0.0143 | 0.935 |
| zinc finger | 0.000907 | 0.000338 | 0.999 |

Lastly, m6A modification is an abundant RNA modification that affects RNA structure [48]. Since m6A modifications affect RNA folding but are not considered in currently-available *in silico* folding algorithms, we tested whether m6A-iCLIP (miCLIP-seq) data could be used as an input for nearBynding to observe protein binding contexts relative to m6A modification. Multiple RBPs are involved in the writing, reading, and erasing of m6A, such as RBM15, IGF2BP1/3, and FTO, respectively [49–51]. We used miCLIP-seq data [52] and eCLIP data [28] from HepG2 to determine whether these m6A-interacting RBPs show binding preferences relative to m6A modification. As expected, RBM15, IGF2BP1, and IGF2BP3 all demonstrated a preference for binding m6A-modified RNA (Fig. 5B). In contrast, FTO did not seem to occupy m6A-modified regions of the transcriptome, perhaps reflecting its role as an m6A eraser. Though the density amplitude for these analyses were modest, likely due to a small signal to noise ratio in the miCLIP-seq data, they demonstrate the diversity of data types that can inform RBP binding contexts using nearBynding.

## Discussion

nearBynding provides a pipeline to discern RNA structure at and proximal to the site of protein binding within regions of the transcriptome defined by the user. Protein-binding data can be input as either aligned CLIP reads or peak-called files. RNA structure can be input based on *in silico* sequence-based prediction or derived from experiments. RNA structure binding profiles can be visually and quantitatively compared across multiple formats. Using simulation datasets, we verified that nearBynding is capable of discerning signal even with a foreground to background ratio of 0.05 and peak concordance less than 40%. Using extensive ENCODE eCLIP datasets, we established that nearBynding reproduces binding profiles between biological replicates and across cell lines. In addition, these binding profiles recapitulated known RBP binding preferences and identified protein domains that may bind G4.

Our analyses further revealed that some RBPs demonstrate cell-type-specific binding profiles. The same RBP may have dissimilar binding profiles between cell types due to different target transcripts being expressed, other RBPs affecting RBP targets, or different co-binders affecting the binding of the interrogated RBP between cell types. Therefore, RBP binding profiles should not be assumed based on existing data; instead, RBP binding data for the interrogated cell type ought to be used if available. Future comprehensive analyses using multiple cell lines may reveal consensus binding profiles for a subset of RBPs or among groups of cell types.

Of note, a strong cross-correlation signal between RBP binding and RNA structures does not necessarily imply that an RBP binds that specific structure. It is possible that the RNA is prone to adopting a structure when it is not bound by the RBP. DROSHA, for example, binds G4-forming regions only when these regions are unfolded [43]. Because many G4-forming sequences are actively unfolded *in vivo*, we cannot know without further molecular experimentation whether an RBP binds to G4s or RBP-associated sequences are prone to forming G4s. We speculate that a phenomenon similar to DROSHA’s binding drove the enrichment of dsRBD-containing RBPs among the higher G4 signals (Table 1), since G4-forming sequences are necessarily GC-rich and likely form stable regions of dsRNA. Biochemical experimentation such as kinetics assays or crystal structures are necessary to definitively ascertain RBP binding.

One limitation of nearBynding is that it does not easily accommodate *in vitro* RBP binding data such as those derived from SELEX or RNA Bind-N-Seq. To incorporate these in vitro data, the user would need to create an annotated genome containing sequences for every oligonucleotide probe in the queried experiment. Currently, the nearBynding pipeline uses aligned CLIP-seq data to evaluate RBP binding preferences, but it does not require peak-calling. It has been previously observed that modeling RBP binding as a list of bound regions across the transcriptome provides only a coarse approximation of RBP binding motifs [19]. Quantitative RBP affinities for different segments of the transcriptome, such as is provided by the amplitudes of aligned CLIP-seq reads, may be better used to distinguish RBP target motifs according to their binding frequency. Some differences may exist in RBP binding profiles when CLIPper-called peaks, Piranha-called peaks, or aligned eCLIP reads were served as the input for the same experiment. However, the differences between binding profiles for different RBPs is far greater than the differences for the same RBP interrogated using different inputs, demonstrating the possibility of omitting the step of peak calling for RBP binding analysis. Additionally, nearBynding currently only supports the consideration of background signal for RBP binding, such as for an input control in a CLIP experiment. Future work will support the possibility of removing background RNA structure signal, such as for an input control in RNA immunoprecipitation (RIP) experiments that use antibodies targeting RNA structures or modifications [51,53].

Notably, nearBynding does not incorporate RNA sequence into the RBP binding profiles as a default. While sequence is inarguably an important component of RBP binding, our primary aim is to elaborate on how structure-based folding predictions are addressed. However, in cases of single-nucleotide RBP binding information such as iCLIP or hiCLIP [33,54] (see Fig. 4D), it would likely be possible to assess RNA sequence preferences relative to RBP binding by separating each of the four nucleotides into individual RNA tracks, similar to how CapR separates six RNA structures into different tracks.

To our knowledge, every state-of-the-art algorithm that incorporates RNA structure into predictions of RBP binding motifs relies on RNA sequence alone to predict RNA secondary structure [55]. Similarly, all nearBynding analyses that use CapR-predicted RNA structures assume that the mRNA being folded is naked and unmodified, with only the queried RBP binding it. To address these limitations, we encoded the option for users to input experimentally-derived RNA structure information. This flexibility could be used to study the binding of non-canonical RNA structures (e.g., G4s, triple helices) and RNA modifications, such as m6A or N^4^-acetylcytidine. In addition, users can improve the study of canonical RNA structure binding by incorporating structural information collected via chemical probing (e.g., selective 2’-hydroxyl acylation analyzed by primer extension [SHAPE] or dimethyl sulfate [DMS]). Future work will characterize the impact of chemical probing-informed RNA structure data on RBP profiles relative to *in silico*-derived RNA structure.

Our long-term goal in developing nearBynding is to interrogate RBP binding relative to RNA structure, but users could alternatively study the binding of one RBP relative to another instead. In this case, we will input both RBPs’ binding data as individual tracks for analyses. Future work will build on this idea to characterize RBP complexes relative to RNA structure.

### Availability of data and materials

The nearBynding pipeline is available at Bioconductor (https://bioconductor.org/packages/nearBynding/, BioC 3.12). nearBynding v99.12 was used for all data analysis and the latest updates are at https://github.com/vbusa1/nearBynding. nearBynding is entirely coded in R (v4.0) and its license is Artistic-2.0. nearBynding was run on macOS Catalina (10.15.4) but has been effectively run on Windows and Linux OS as well. Some processed data and the code necessary to generate the simulated data and figures presented in this paper are available at https://github.com/vbusa1/nearBynding_manuscript. CapR [6] (v1.1.1) is available at https://github.com/fukunagatsu/CapR and StereoGene [26] (v2.20) is available at http://stereogene.bioinf.fbb.msu.ru/.

## Methods

### nearBynding inputs

In order to provide predicted RNA structure context for RBP binding, the nearBynding pipeline requires the following pieces of input data: (1) CLIP-seq alignment tracks for the RBP of interest, (2) an annotated genome and associated FASTA sequence, and (3) a list of transcripts of interest. It is recommended that all transcripts selected are expressed in the cell type used for the CLIP-seq experiment. Alternative RNA structure information can optionally be included, and it is recommended that the data is derived from the same cell type.

#### RBP binding data

All eCLIP BAM files and CLIPper peak files are available in the ENCODE repository (v102 https://www.encodeproject.org/) [28]. STAU1 hiCLIP data is available in the iMaps repository (https://imaps.genialis.com/iclip/search/collection/hi-clip-reveals-m-rna-secondary-structures) [33]. UPF1 WT, K498A, and DEAA CLIP-seq data is from [34] (GEO: GSE69586). Peak calling was conducted using Piranha [36] v1.2.1 (http://smithlabresearch.org/software/piranha/) as described in the original paper: bin size of 36 nucleotides and the single covariate being the log of the read counts from the control input.

#### Genome

We used FASTA and GTF files from the Ensembl release 100 Homo sapiens GRCh38 genome (ftp://ftp.ensembl.org/pub/release-100/) [56].

#### List of transcripts

HepG2 and K562 isoform information was derived from RNA-seq data available in the ENCODE repository (identifiers ENCSR181ZGR and ENCSR885DVH, respectively); HEK293 isoform information was derived from WT RNA-seq data [57] (accessible through the Gene Expression Omnibus (GEO) accession number GSE122425).

#### Experimental RNA structure data

Single-nucleotide resolution profiling of m6A (miCLIP-seq) in HepG2 cells is from [52] (GEO: GSE121942). rG4-seq data in HEK293 cells is from [39] (GEO: GSE77282). Both of these datasets were aligned to hg19 and so were lifted over to hg38 using a UCSC chain file before input into nearBynding (http://hgdownload.soe.ucsc.edu/goldenPath/hg19/liftOver/hg19ToHg38.over.chain.gz) [58].

### Map data to pseudochromosomes

Users must first choose which regions of the transcripts of interest to interrogate (e.g. UTRs, exons, whole transcript), based on annotations available in the genome GTF file. nearBynding creates (1) a chain file that will be used to map the selected regions of transcripts end-to-end, excluding the intergenic regions and undesired transcripts that comprise the majority of the genome, and (2) a file detailing the names and sizes of all the chain file’s pseudochromosomes. The chain file can then be used to transfer genome references of the CLIP-seq data from the whole-genome alignment to the transcriptome alignment of interest. If external RNA structure data is being studied, its genome references would need to be transferred to the transcriptome alignment as well using the same chain file. Chain files cannot accommodate overlapping intervals since they cause ambiguous regions in the transfer process, so it is recommended that users supply the highest-expressed isoform of every gene expressed in the cell type of the CLIP-seq data to create the concatenated pseudochromosomes.

### CapR RNA structure prediction

nearBynding pulls the sequences of selected regions of transcripts of interest from the genome FASTA file based on genome annotations using bedtools [59] (https://bedtools.readthedocs.io/en/latest/). Probabilistic RNA structure for the selected regions are derived from *in silico* folding predictions by CapR, which includes RNAfold software in its structure predictions [6]. Each nucleotide is scored as having a 0 to 1 probability of adopting one of six different contexts by CapR. The data for the six different folding conformations are then separated and the transcript fragments are concatenated into pseudo-chromosomes. In secondary structure representation, RNA bases are depicted as vertices of polygons with edges of RNA backbone or hydrogen bonds (Fig. 1B). The six different RNA structure contexts are defined thus: stem context is if a base participates in hydrogen-bonding with another base; exterior context is if a base does not form a polygon such as that the end of a transcript; hairpin context is if a single-stranded base is involved in a polygon with only one hydrogen-bonding edge; bulge context is if a single-stranded base is involved in a polygon with two hydrogen-bonding edges and where all stem context vertices are contiguous in the polygon; internal context is if a single-stranded base is involved in a polygon with two hydrogen-bonding edges and where stem context vertices are not contiguous in the polygon; multibranch context is if a single-stranded base is involved in a polygon with at least three hydrogen-bonding edges [6].

### Relative binding position calculation

To visualize the RNA structure landscape surrounding protein binding, StereoGene [26] is used to calculate the cross-correlation between RNA structure and protein binding. nearBynding analyzes local structure in single-nucleotide frames, which sacrifices some of the computational efficiency of StereoGene but maximizes RBP binding resolution. Cross-correlation densities are within the range −1 to +1, where −1 suggests perfect depletion of an RBP for a tested RNA structure context, 0 represents no binding preference, and +1 suggests perfect RBP binding for a tested RNA structure context. Since actual correlation densities are far more modest, they are reported as density x 100 for visualization.

### nearBynding output analyses

nearBynding allows users to calculate the similarity between output binding profiles via Wasserstein distance, where small values indicate greater similarity. Users can also assess the grouping of different categories of points via bootstrapping and the Kolmogorov–Smirnov test (Fig. S5C).

The protein domain data used to compare G4 binding preferences is available in the Pfam repository [46] (v33.1 http://pfam.xfam.org/). RGG-repeat-containing proteins are listed in the supplementary data of ref. [60].

## Declarations

### Ethics approval and consent to participate

Not applicable.

### Consent for publication

Not applicable.

### Authors’ contributions

All authors were involved in conceptualization of the project. V.F.B. wrote the pipeline and ran all bioinformatic analyses. A.V.F. contributed coding suggestions to and reviewed the pipeline. V.F.B wrote the original draft; all authors reviewed and edited the manuscript. A.V.F., A.K.L.L., and E.J.F. provided supervision and guidance throughout the project. All authors read and approved the final manuscript.

## Acknowledgements

We thank Leung lab members, Fertig lab members, and Tim Nieuwenhuis for crowd-sourced validation and ongoing suggestions for the project; Kurt Weir for discussion and reading the manuscript; Arjun Bhutkar for the suggestion to use Wasserstein distances; and Eric Van Nostrand for providing processed data from ref. [30].

---

**Figure S1.**
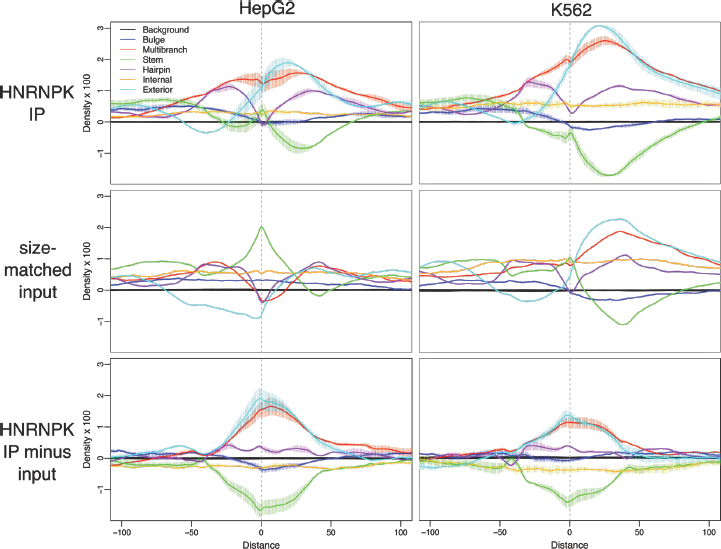
Binding profiles of HNRNPK immunoprecipitation alone (top), size-matched input alone (middle), or HNRNPK IP minus size-matched input signal (bottom) for HepG2 (left) and K562 (right) cell lines.

**Figure S2.**
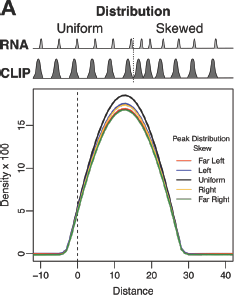
Graphical depiction of simulated skewing and cross-correlation distribution tracks of simulated data with differing peak skews.

**Figure S3.**
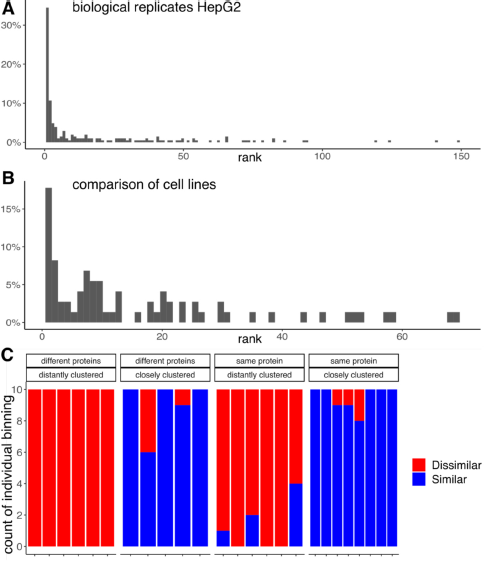
**A.** Histogram of Wasserstein distance ranks of paired biological replicates in HepG2 cells. **B.** Histogram of Wasserstein distance ranks for the same RBP in HepG2 versus K562 cell lines. **C.** Proportion of each pairwise comparison binned either similar (blue) or dissimilar (red) by crowd-sourced validation and organized by Wasserstein distance-determined similarity.

**Figure S4.**
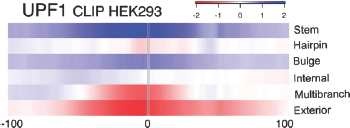
Binding profile of helicase-dependent UPF1 binding based on subtraction of K498A UPF1 signal from WT.

**Figure S5.**
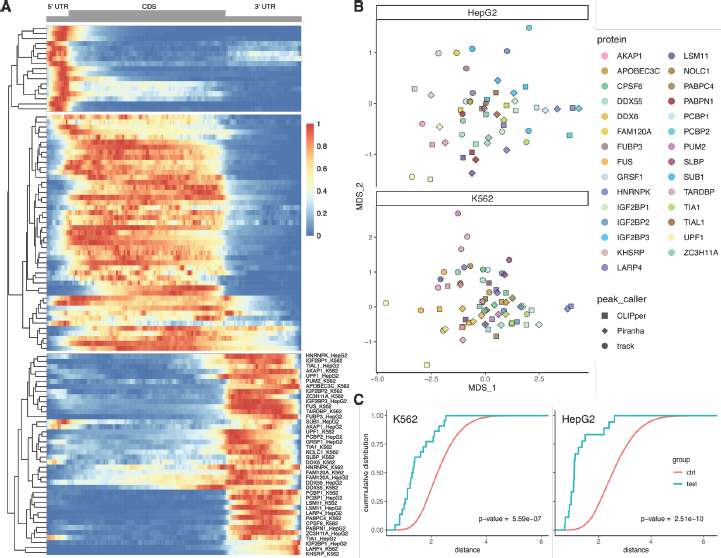
**A.** Heat map of high-signal eCLIP samples across mRNAs, generated as in ref. [30].RBP and cell type are listed for RBPs that primarily bind 3’UTRs. **B.** Distances between binding profiles with RBP binding defined by peak-callers or aligned reads mapped into Cartesian space via multidimensional scaling. Signal was tested across all six predicted RNA structure contexts in HepG2 (top) and K562 (bottom) cell lines. **C.** Kolmogorov-Smirnov test comparing the mean distance between binding profiles for Piranha, CLIPper, and aligned-read inputs of the same protein versus bootstrapping for three random binding profiles in K562 (left) and HepG2 (right) cell lines.

**Figure S6.**
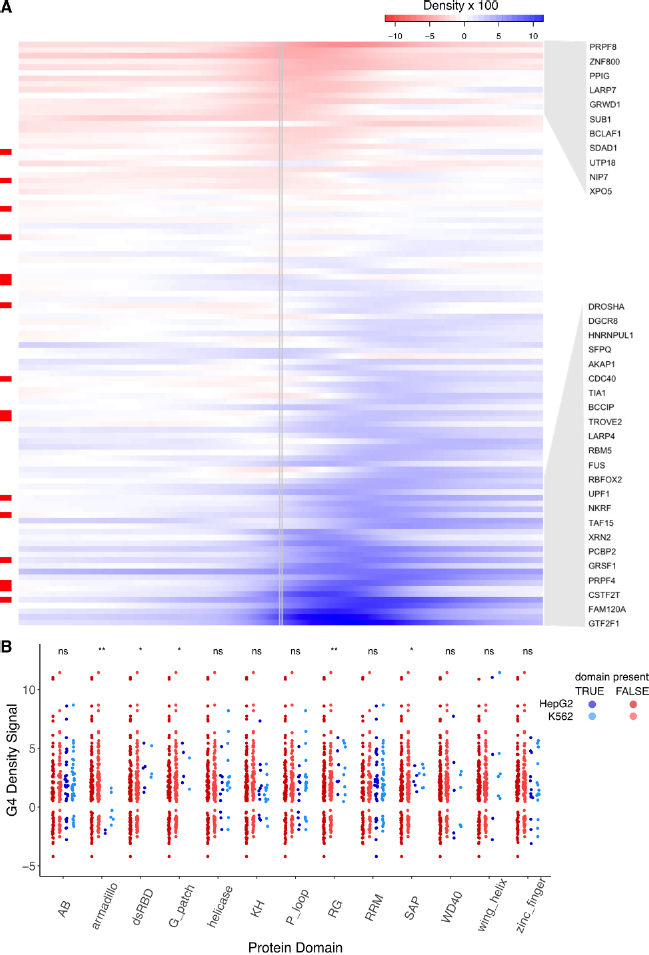
**A.** G4 binding profiles for all 103 proteins with eCLIP data in HepG2 cells. RBPs with molecular evidence of G4 binding in the literature are indicated in red along the left. RBPs with the most positive and most negative correlation signals are highlighted by grey blocks and listed on the right. **B.** Sina plots comparing maximal G4 binding signals that contain or do not contain the protein domains indicated for K562 and HepG2 proteins. Protein domains tested are alpha-beta (AB), armadillo, double-stranded RNA-binding (dsRBD), G-patch, helicase, K homology (KH), phosphate-binding loop (P-loop), RGG-repeats (RG), RNA-recognition motif (RRM), SAP, WD40, winged helix, and zinc finger.

## References

1. Gerstberger S, Hafner M, Tuschl T. A census of human RNA-binding proteins. Nature Reviews Genetics [Internet]. 2014 [cited 2019 Aug 29];15:829–45. Available from: https://www.nature.com/articles/nrg3813

2. Wang Z-L, Li B, Luo Y-X, Lin Q, Liu S-R, Zhang X-Q, et al. Comprehensive Genomic Characterization of RNA-Binding Proteins across Human Cancers. Cell Reports [Internet]. 2018 [2019 Aug 20];22:286–98. Available from: http://www.sciencedirect.com/science/article/pii/S2211124717318442

3. Fredericks AM, Cygan KJ, Brown BA, Fairbrother WG. RNA-Binding Proteins: Splicing Factors and Disease. Biomolecules [Internet]. 2015 [cited 2019 Aug 20];5:893–909. Available from: https://www.ncbi.nlm.nih.gov/pmc/articles/PMC4496701/

4. Ye B, Petritsch C, Clark IE, Gavis ER, Jan LY, Jan YN. nanos and pumilio Are Essential for Dendrite Morphogenesis in Drosophila Peripheral Neurons. Current Biology. 2004;14:314–21.

5. Kazan H, Ray D, Chan ET, Hughes TR, Morris Q. RNAcontext: A New Method for Learning the Sequence and Structure Binding Preferences of RNA-Binding Proteins. PLoS Comput Biol [Internet]. 2010 [cited 2019 Jul 11];6. Available from: https://www.ncbi.nlm.nih.gov/pmc/articles/PMC2895634/

6. Fukunaga T, Ozaki H, Terai G, Asai K, Iwasaki W, Kiryu H. CapR: revealing structural specificities of RNA-binding protein target recognition using CLIP-seq data. Genome Biol. 2014;15:R16.

7. Arguello AE, DeLiberto AN, Kleiner RE. RNA Chemical Proteomics Reveals the N6-Methyladenosine (m6A)-Regulated Protein–RNA Interactome. J Am Chem Soc [Internet]. 2017 [cited 2020 Feb 20];139:17249–52. Available from: https://doi.org/10.1021/jacs.7b09213

8. Heller D, Krestel R, Ohler U, Vingron M, Marsico A. ssHMM: extracting intuitive sequence-structure motifs from high-throughput RNA-binding protein data. Nucleic Acids Res. 2017;45:11004–18.

9. Blaszczyk J, Gan J, Tropea JE, Court DL, Waugh DS, Ji X. Noncatalytic Assembly of Ribonuclease III with Double-Stranded RNA. Structure. 2004;12:457–66.

10. Li X, Quon G, Lipshitz HD, Morris Q. Predicting in vivo binding sites of RNA-binding proteins using mRNA secondary structure. RNA. 2010;16:1096–107.

11. Bahrami-Samani E, Penalva LOF, Smith AD, Uren PJ. Leveraging cross-link modification events in CLIP-seq for motif discovery. Nucleic Acids Res. 2015;43:95–103.

12. Edupuganti RR, Geiger S, Lindeboom RGH, Shi H, Hsu PJ, Lu Z, et al. N6-methyladenosine (m6A) recruits and repels proteins to regulate mRNA homeostasis. Nat Struct Mol Biol. 2017;24:870–8.

13. Wilbert ML, Huelga SC, Kapeli K, Stark TJ, Liang TY, Chen SX, et al. LIN28 binds messenger RNAs at GGAGA motifs and regulates splicing factor abundance. Mol Cell. 2012;48:195–206.

14. Taliaferro JM, Lambert NJ, Sudmant PH, Dominguez D, Merkin JJ, Alexis MS, et al. RNA sequence context effects measured in vitro predict in vivo protein binding and regulation. Mol Cell [Internet]. 2016 [cited 2019 Jul 12];64:294–306. Available from: https://www.ncbi.nlm.nih.gov/pmc/articles/PMC5107313/

15. Li X, Kazan H, Lipshitz HD, Morris QD. Finding the target sites of RNA-binding proteins. Wiley Interdiscip Rev RNA [Internet]. 2014 [cited 2019 Aug 6];5:111–30. Available from: https://www.ncbi.nlm.nih.gov/pmc/articles/PMC4253089/

16. Lorenz R, Bernhart SH, Höner zu Siederdissen C, Tafer H, Flamm C, Stadler PF, et al. ViennaRNA Package 2.0. Algorithms Mol Biol [Internet]. 2011 [cited 2018 Sep 18];6:26. Available from: https://www.ncbi.nlm.nih.gov/pmc/articles/PMC3319429/

17. Ding Y, Lawrence CE. A statistical sampling algorithm for RNA secondary structure prediction. Nucleic Acids Res [Internet]. 2003 [cited 2019 Jul 12];31:7280–301. Available from: https://www.ncbi.nlm.nih.gov/pmc/articles/PMC297010/

18. Steffen P, Voß B, Rehmsmeier M, Reeder J, Giegerich R. RNAshapes: an integrated RNA analysis package based on abstract shapes. Bioinformatics [Internet]. 2006 [cited 2019 Jul 16];22:500–3. Available from: https://academic.oup.com/bioinformatics/article/22/4/500/184565

19. Maticzka D, Lange SJ, Costa F, Backofen R. GraphProt: modeling binding preferences of RNA-binding proteins. Genome Biol [Internet]. 2014 [cited 2019 Jul 11];15:R17. Available from: https://www.ncbi.nlm.nih.gov/pmc/articles/PMC4053806/

20. Pan X, Shen H-B. Predicting RNA–protein binding sites and motifs through combining local and global deep convolutional neural networks. Bioinformatics [Internet]. 2018 [cited 2019 Jul 12];34:3427–36. Available from: https://academic.oup.com/bioinformatics/article/34/20/3427/4990826

21. Yan Z, Hamilton WL, Blanchette M. Graph neural representational learning of RNA secondary structures for predicting RNA-protein interactions. Bioinformatics [Internet]. 2020 [cited 2020 Jul 25];36:i276–84. Available from: https://www.ncbi.nlm.nih.gov/pmc/articles/PMC7355240/

22. Chen X, Castro SA, Liu Q, Hu W, Zhang S. Practical considerations on performing and analyzing CLIP-seq experiments to identify transcriptomic-wide RNA-protein interactions. Methods [Internet]. 2019 [cited 2020 Feb 14];155:49–57. Available from: http://www.sciencedirect.com/science/article/pii/S1046202318302317

23. Sasse A, Laverty KU, Hughes TR, Morris QD. Motif models for RNA-binding proteins. Current Opinion in Structural Biology [Internet]. 2018 [cited 2019 Aug 13];53:115–23. Available from: http://www.sciencedirect.com/science/article/pii/S0959440X18300654

24. Carlile TM, Martinez NM, Schaening C, Su A, Bell TA, Zinshteyn B, et al. mRNA structure determines modification by pseudouridine synthase 1. Nat Chem Biol [Internet]. 2019 [cited 2019 Sep 6];1–9. Available from: https://www.nature.com/articles/s41589-019-0353-z

25. Jarmoskaite I, Denny SK, Vaidyanathan PP, Becker WR, Andreasson JOL, Layton CJ, et al. A Quantitative and Predictive Model for RNA Binding by Human Pumilio Proteins. Molecular Cell [Internet]. 2019 [cited 2019 Dec 9];74:966-981.e18. Available from: http://www.sciencedirect.com/science/article/pii/S1097276519302825

26. Stavrovskaya ED, Niranjan T, Fertig EJ, Wheelan SJ, Favorov AV, Mironov AA. StereoGene: rapid estimation of genome-wide correlation of continuous or interval feature data. Bioinformatics [Internet]. 2017 [cited 2018 Oct 9];33:3158–65. Available from: http://academic.oup.com/bioinformatics/article/33/20/3158/3957591

27. Schuhmacher D, diagrams) BB (aha and power, Bonneel (networkflow) N, shortlist) CG (simplex and, Hartmann (semidiscrete1) V, integration) FH (transport_track and networkflow, et al. transport: Computation of Optimal Transport Plans and Wasserstein Distances [Internet]. 2020 [cited 2020 Apr 21]. Available from: https://CRAN.R-project.org/package=transport

28. Davis CA, Hitz BC, Sloan CA, Chan ET, Davidson JM, Gabdank I, et al. The Encyclopedia of DNA elements (ENCODE): data portal update. Nucleic Acids Res. 2018;46:D794–801.

29. Beaudoin J-D, Jodoin R, Perreault J-P. New scoring system to identify RNA G-quadruplex folding. Nucleic Acids Res [Internet]. 2014 [cited 2020 Jul 21];42:1209–23. Available from: https://www.ncbi.nlm.nih.gov/pmc/articles/PMC3902908/

30. Van Nostrand EL, Pratt GA, Yee BA, Wheeler EC, Blue SM, Mueller J, et al. Principles of RNA processing from analysis of enhanced CLIP maps for 150 RNA binding proteins. Genome Biology [Internet]. 2020 [cited 2020 Apr 11];21:90. Available from: https://doi.org/10.1186/s13059-020-01982-9

31. Nam Y, Chen C, Gregory RI, Chou JJ, Sliz P. Molecular Basis for Interaction of let-7 MicroRNAs with Lin28. Cell [Internet]. 2011 [cited 2020 May 3];147:1080–91. Available from: http://www.sciencedirect.com/science/article/pii/S0092867411012669

32. Ramos A, Grünert S, Adams J, Micklem DR, Proctor MR, Freund S, et al. RNA recognition by a Staufen double-stranded RNA-binding domain. EMBO J [Internet]. 2000 [cited 2019 Aug 15];19:997–1009. Available from: https://www.ncbi.nlm.nih.gov/pmc/articles/PMC305639/

33. Sugimoto Y, Vigilante A, Darbo E, Zirra A, Militti C, D’Ambrogio A, et al. hiCLIP reveals the in vivo atlas of mRNA secondary structures recognized by Staufen 1. Nature [Internet]. 2015 [cited 2019 Oct 16];519:491–4. Available from: https://www.ncbi.nlm.nih.gov/pmc/articles/PMC4376666/

34. Lee SR, Pratt G, Martinez F, Yeo GW, Lykke-Andersen J. Target discrimination in nonsense-mediated mRNA decay requires Upf1 ATPase activity. Mol Cell [Internet]. 2015 [cited 2018 Sep 20];59:413–25. Available from: https://www.ncbi.nlm.nih.gov/pmc/articles/PMC4673969/

35. Lovci MT, Ghanem D, Marr H, Arnold J, Gee S, Parra M, et al. Rbfox proteins regulate alternative mRNA splicing through evolutionarily conserved RNA bridges. Nat Struct Mol Biol [Internet]. 2013 [cited 2020 Apr 29];20:1434–42. Available from: https://www.ncbi.nlm.nih.gov/pmc/articles/PMC3918504/

36. Uren PJ, Bahrami-Samani E, Burns SC, Qiao M, Karginov FV, Hodges E, et al. Site identification in high-throughput RNA–protein interaction data. Bioinformatics [Internet]. 2012 [cited 2020 Apr 28];28:3013–20. Available from: https://www.ncbi.nlm.nih.gov/pmc/articles/PMC3509493/

37. Brázda V, Hároníková L, Liao JCC, Fojta M. DNA and RNA Quadruplex-Binding Proteins. Int J Mol Sci [Internet]. 2014 [cited 2019 Dec 16];15:17493–517. Available from: https://www.ncbi.nlm.nih.gov/pmc/articles/PMC4227175/

38. Puig Lombardi E, Londoño-Vallejo A. A guide to computational methods for G-quadruplex prediction. Nucleic Acids Res [Internet]. 2020 [cited 2020 Jul 21];48:1–15. Available from: https://www.ncbi.nlm.nih.gov/pmc/articles/PMC6943126/

39. Kwok CK, Marsico G, Sahakyan AB, Chambers VS, Balasubramanian S. rG4-seq reveals widespread formation of G-quadruplex structures in the human transcriptome. Nature Methods [Internet]. 2016 [cited 2019 Nov 1];13:841–4. Available from: https://www.nature.com/articles/nmeth.3965

40. Simko EAJ, Liu H, Zhang T, Velasquez A, Teli S, Haeusler AR, et al. G-quadruplexes offer a conserved structural motif for NONO recruitment to NEAT1 architectural lncRNA. Nucleic Acids Res. 2020;

41. Yagi R, Miyazaki T, Oyoshi T. G-quadruplex binding ability of TLS/FUS depends on the β-spiral structure of the RGG domain. Nucleic Acids Res [Internet]. 2018 [cited 2020 May 5];46:5894–901. Available from: https://www.ncbi.nlm.nih.gov/pmc/articles/PMC6159513/

42. Pietras Z, Wojcik MA, Borowski LS, Szewczyk M, Kulinski TM, Cysewski D, et al. Dedicated surveillance mechanism controls G-quadruplex forming non-coding RNAs in human mitochondria. Nat Commun [Internet]. 2018 [cited 2020 Jul 22];9. Available from: https://www.ncbi.nlm.nih.gov/pmc/articles/PMC6028389/

43. Rouleau SG, Garant J-M, Bolduc F, Bisaillon M, Perreault J-P. G-Quadruplexes influence pri-microRNA processing. RNA Biol [Internet]. 2017 [cited 2020 Jul 22];15:198–206. Available from: https://www.ncbi.nlm.nih.gov/pmc/articles/PMC5798953/

44. Zyner KG, Mulhearn DS, Adhikari S, Martínez Cuesta S, Di Antonio M, Erard N, et al. Genetic interactions of G-quadruplexes in humans. eLife [Internet]. [cited 2020 Jul 22];8. Available from: https://www.ncbi.nlm.nih.gov/pmc/articles/PMC6615864/

45. Lee DSM, Ghanem LR, Barash Y. Integrative analysis reveals RNA G-quadruplexes in UTRs are selectively constrained and enriched for functional associations. Nature Communications [Internet]. 2020 [cited 2020 Jan 28];11:1–12. Available from: https://www.nature.com/articles/s41467-020-14404-y

46. El-Gebali S, Mistry J, Bateman A, Eddy SR, Luciani A, Potter SC, et al. The Pfam protein families database in 2019. Nucleic Acids Res [Internet]. Oxford Academic; 2019 [cited 2020 Jul 22];47:D427–32. Available from: https://academic.oup.com/nar/article/47/D1/D427/5144153

47. Brázda V, Červeň J, Bartas M, Mikysková N, Coufal J, Pečinka P. The Amino Acid Composition of Quadruplex Binding Proteins Reveals a Shared Motif and Predicts New Potential Quadruplex Interactors. Molecules [Internet]. 2018 [cited 2020 May 5];23. Available from: https://www.ncbi.nlm.nih.gov/pmc/articles/PMC6225207/

48. Roost C, Lynch SR, Batista PJ, Qu K, Chang HY, Kool ET. Structure and Thermodynamics of N6-Methyladenosine in RNA: A Spring-Loaded Base Modification. J Am Chem Soc [Internet]. 2015 [cited 2020 Oct 5];137:2107–15. Available from: https://www.ncbi.nlm.nih.gov/pmc/articles/PMC4405242/

49. Yu J, Chen M, Huang H, Zhu J, Song H, Zhu J, et al. Dynamic m6A modification regulates local translation of mRNA in axons. Nucleic Acids Res [Internet]. 2018 [cited 2020 Jul 23];46:1412–23. Available from: https://www.ncbi.nlm.nih.gov/pmc/articles/PMC5815124/

50. Patil DP, Chen C-K, Pickering BF, Chow A, Jackson C, Guttman M, et al. m6A RNA methylation promotes XIST-mediated transcriptional repression. Nature [Internet]. 2016 [cited 2020 Jul 23];537:369–73. Available from: https://www.ncbi.nlm.nih.gov/pmc/articles/PMC5509218/

51. Huang H, Weng H, Sun W, Qin X, Shi H, Wu H, et al. Recognition of RNA N6-methyladenosine by IGF2BP Proteins Enhances mRNA Stability and Translation. Nat Cell Biol [Internet]. 2018 [cited 2020 Feb 18];20:285–95. Available from: https://www.ncbi.nlm.nih.gov/pmc/articles/PMC5826585/

52. Huang H, Weng H, Zhou K, Wu T, Zhao BS, Sun M, et al. Histone H3 trimethylation at lysine 36 guides m6A RNA modification co-transcriptionally. Nature [Internet]. 2019 [cited 2020 Apr 12];567:414–9. Available from: https://www.ncbi.nlm.nih.gov/pmc/articles/PMC6438714/

53. Park OH, Ha H, Lee Y, Boo SH, Kwon DH, Song HK, et al. Endoribonucleolytic Cleavage of m6A-Containing RNAs by RNase P/MRP Complex. Molecular Cell [Internet]. 2019 [cited 2019 Oct 29];74:494-507.e8. Available from: http://www.sciencedirect.com/science/article/pii/S1097276519301455

54. Huppertz I, Attig J, D’Ambrogio A, Easton LE, Sibley CR, Sugimoto Y, et al. iCLIP: Protein–RNA interactions at nucleotide resolution. Methods [Internet]. 2014 [cited 2020 Mar 11];65:274–87. Available from: https://www.ncbi.nlm.nih.gov/pmc/articles/PMC3988997/

55. Guo JU, Bartel DP. RNA G-quadruplexes are globally unfolded in eukaryotic cells and depleted in bacteria. Science [Internet]. 2016 [cited 2019 Nov 1];353. Available from: https://www.ncbi.nlm.nih.gov/pmc/articles/PMC5367264/

56. Howe KL, Contreras-Moreira B, De Silva N, Maslen G, Akanni W, Allen J, et al. Ensembl Genomes 2020—enabling non-vertebrate genomic research. Nucleic Acids Res [Internet]. Oxford Academic; 2020 [cited 2020 Jul 26];48:D689–95. Available from: https://academic.oup.com/nar/article/48/D1/D689/5584694

57. Sun Z, Xue S, Xu H, Hu X, Chen S, Yang Z, et al. Effects of NSUN2 deficiency on the mRNA 5-methylcytosine modification and gene expression profile in HEK293 cells. Epigenomics [Internet]. 2018 [cited 2020 Feb 22];11:439–53. Available from: https://www.futuremedicine.com/doi/full/10.2217/epi-2018-0169

58. Lee CM, Barber GP, Casper J, Clawson H, Diekhans M, Gonzalez JN, et al. UCSC Genome Browser enters 20th year. Nucleic Acids Res [Internet]. 2020 [cited 2020 Jul 26];48:D756–61. Available from: https://www.ncbi.nlm.nih.gov/pmc/articles/PMC7145642/

59. Quinlan AR. BEDTools: the Swiss-army tool for genome feature analysis. Curr Protoc Bioinformatics [Internet]. 2014 [cited 2020 Jul 26];47:11.12.1-11.12.34. Available from: https://www.ncbi.nlm.nih.gov/pmc/articles/PMC4213956/

60. Ozdilek BA, Thompson VF, Ahmed NS, White CI, Batey RT, Schwartz JC. Intrinsically disordered RGG/RG domains mediate degenerate specificity in RNA binding. Nucleic Acids Res [Internet]. Oxford Academic; 2017 [cited 2020 Jun 18];45:7984–96. Available from: https://academic.oup.com/nar/article/45/13/7984/3855590

